# Oxidative-stress related increase in keratoconus tear MDA and GPX3 while NRF2-antioxidant functions decrease in stromal cells

**DOI:** 10.1101/2025.11.13.688300

**Authors:** Madhuri A Koduri, Mackenzie Charter, Rohini Sonar, Rashmi Deshmukh, Christina R Prescott, Rose Mandel, Lawrence Sperber, Ting-Fang Lee, Elias H Kahan, Ilyse D Haberman, Vivek Singh, Andrea L Blitzer, George Maiti, Shukti Chakravarti

## Abstract

Keratoconus (KC) is a common eye disease where the cornea undergoes degenerative thinning and steepening. The absence of biomarkers for early diagnosis prior to the onset of overt corneal phenotypes and the lack of curative treatments rooted in a fundamental understanding of KC biology remain significant challenges. To address these issues, we investigated the role of unresolved oxidative stress in KC pathogenesis. Malondialdehyde (MDA) a lipid peroxidation byproduct that accumulates during oxidative stress was significantly elevated in the tears of KC patients compared to unaffected controls and positively correlated with maximal keratometry (Kmax), a measure of KC severity. Similarly, the secreted antioxidant glutathione peroxidase 3 (GPX3), was significantly increased in patient tears, and strongly correlated with Kmax. In a cell culture model of oxidative stress, KC corneal stromal cells displayed increased apoptosis and suboptimal activation of NRF2, a transcription factor master regulator of antioxidant genes. Conversely, inhibition of NRF2 in donor stromal cells elicited KC-like cellular phenotype, whereas sulforaphane, an NRF2 booster restored antioxidant gene expression and the deposition of cornea-typical collagens. Our study identified cellular antioxidant signaling dysregulations in keratoconus where sulforaphane treatment may be restorative. Consistent increases in patient tear MDA and GPX3 present these as promising biomarkers for KC diagnosis and severity predictions.

## Introduction

Keratoconus (KC) is a common eye disease characterized by the progressive thinning and weakening of the cornea (1–4), leading to irregular astigmatism, scarring, and vision loss. At the corneal tissue level there are reports on abnormal epithelial thickening, loss of stromal keratocytes and extracellular matrix (ECM) (5–10). Typically diagnosed in childhood, KC can progress into middle age, with complications such as corneal scarring and acute hydrops. There are no curative treatments for keratoconus, but the progressive weakening of the cornea is managed by UV mediated crosslinking of the collagen fibrils in the stroma, and vision loss is managed by corrective contact lenses and glasses (11–13). Ultimately, severe cases require corneal transplantation.

Keratoconus is multifactorial; both genetic and interacting environmental factors are implicated in keratoconus (14, 15). Studies of familial and nonsyndromic isolated cases suggest a few hundred risk-enhancing rare genetic variants (16–18). While common genetic variants associated with at-risk phenotypes, such as low central corneal thickness and corneal biomechanical properties increase polygenic risk and disease susceptibility (18–20). Thus, multiple genes and pathways affect cellular functions and the stromal ECM they produce.

Early diagnosis, based on a mechanistic understanding of the disease is needed. The eye is naturally subject to a myriad of environmental stresses such as UV and visible light, air pollution, toxins, hypoxia and extreme temperature that cause oxidative stress. A recurring concept is that this oxidative stress is not resolved in keratoconus (21–23). Using this as a premise, here we investigate the presence of antioxidants and oxidative stress related factors in tear fluid of KC subjects. Second, to determine whether there are fundamental differences in the antioxidant response in the stromal at the cellular level, we extracted keratocytes from discarded KC corneal tissues and control donor corneas, cultured as low passage fibroblasts and tested their response to experimental oxidative stress. While the corneal epithelium is also subject to oxidative stress, we focused on the keratocytes as these produce the stromal ECM and ultimately responsible for the biomechanically competent, vision supportive cornea that is impaired in keratoconus.

Our prior transcriptomic and proteomic studies of KC corneas detected down regulations of many antioxidant enzymes, such as, glutathione S transferase, heme oxygenase and thioredoxin reductase 1 pointing to dysregulations in their upstream regulator, the transcription factor nuclear factor erythroid 2-related factor 2 (*NRF2*) (23–25). *NRF2* controls over 200 antioxidant and detoxification genes. Under homeostatic conditions, NRF2 undergoes ubiquitination and degradation and thereby maintained at low basal levels in the cytoplasm (26). This involves its binding to kelch-like ECH-associated protein 1 (KEAP1), and the KEAP1-CUL3-RBX1 E3 ubiquitin ligase complex mediated proteasomal degradation. Under oxidative stress, accumulating oxidants or electrophiles interact with sulfhydryl side chains of cysteine residues in KEAP1, causing its conformational change and loss of affinity for NRF2. Accumulating NRF2 translocate to the nucleus where it forms a heterodimer with musculoaponeurotic fibrosarcoma (MAF) proteins and binds to antioxidant response elements (AREs) in the promoter regions of its target genes. This activates a robust antioxidant defense program, upregulating glutathione metabolism and detoxification of reactive oxygen species to restore redox homeostasis and protection against oxidative damage. However, a suboptimal antioxidant program leads to the buildup of reactive superoxides, lipid peroxidation and the accumulation of malondialdehyde (MDA) byproducts. Indeed, more than 20 years ago a study reported increased immunoreactivity for MDA in corneal sections of bullous keratopathy, Fuchs dystrophy and keratoconus (27).

We report significant elevations in MDA in tear fluid samples of KC patients. Among antioxidants, we measured GPX3 protein levels in the tear fluid and found it to be elevated with a significant positive correlation with maximal keratometry (Kmax) in patients. Additionally, we investigated NRF2 dysregulations in KC and donor corneal fibroblasts in culture. We compared low passage stromal fibroblast cultures from KC and donor (DN) controls with respect to their response to experimental oxidative stress and found increased cell death and poor NRF2 functions in KC than in DN stromal cells. Sulforaphane (1-Isothiocyanato-4 butane; SFN), naturally enriched in cruciferous plants, binds to sulfhydryl groups in KEAP1 and interferes with its ability to interact with NRF2 (28). Treatment of our cell culture models with sulforaphane resulted in improved NRF2 functions in KC fibroblasts. Conversely, treatment of DN fibroblasts with a small-molecule inhibitor of NRF2 evoked KC-like cellular phenotype and decreased induction of antioxidant genes.

## Results

### Tear fluid sample information

Tear samples were collected from consented control subjects without keratoconus (non-KC) and KC patients who visited the NYU Faculty Group Practice (FGP) for routine eye examination or keratoconus consultations (**Supplementary Table 1**). The non-KC group included glaucoma suspects, those with myopia, dry eye disease, corneal scar, blepharitis, allergic conjunctivitis, or cataract. Non-KC subjects with active eye inflammation or infections were excluded. All syndromic keratoconus cases, such as those with Down Syndrome were excluded as well. The non-KC and KC were similar in age, 40 ± 17 and 37.24 ± 11.98 years, respectively. Among those with KC, 29 individuals were also diagnosed with other eye conditions, such as exotropia, post-LASIK ectasia, optic disc and retinal disorders or glaucoma, Fuchs dystrophy, dry eye disease, corneal scar, blepharitis, graft failure, allergic conjunctivitis, or Cataract (**Supplementary Table 1)**. 28 KC subjects from our study had no other eye-related diagnoses.

### Elevated MDA in tear fluid of KC patients

As the unstimulated tear fluid yield is quite low (5-8 μl per eye), we pooled samples from both eyes. We measured MDA levels by ELISA in 54 KC and 28 non-KC (unaffected control) subjects. We estimated total protein concentration in each sample (**Supplementary Table 2),** we used 5 μg per sample to measure MDA. On average, KC tear samples had significantly higher levels of MDA levels (47.31 ± 25.34 nMol/ml) than non-KC (18.94± 19.11 nMol/ml) (**Figure 1A**). After stratifying patients into mild, moderate and severe, based on Kmax values **(Supplementary Table 3)**, significant differences were observed only between non-KC and moderate and between non-KC and severe KC cases (p<0.001, ANOVA with Tukey post hoc tests). A linear regression analysis indicated a positive correlation (p=0.01) between subject-level MDA values and eye-level Kmax measurements in KC samples (**Figure 1B**), including both eyes and using cluster-robust standard errors to account for within-patient correlation. We also observed a slight positive correlation between MDA levels and patient age (p=0.17; **Figure 1C**) and disease duration (p=0.08; **Figure 1D**), with neither reaching significance. We found no correlation between MDA levels and sex of the subjects (p=0.96; **Figure 1E**). To determine if systemic factors regulate MDA levels, we measured MDA in the plasma of a subset of KC and non-KC controls and found these to be comparable (**Figure 1F**).

**Figure 1:**
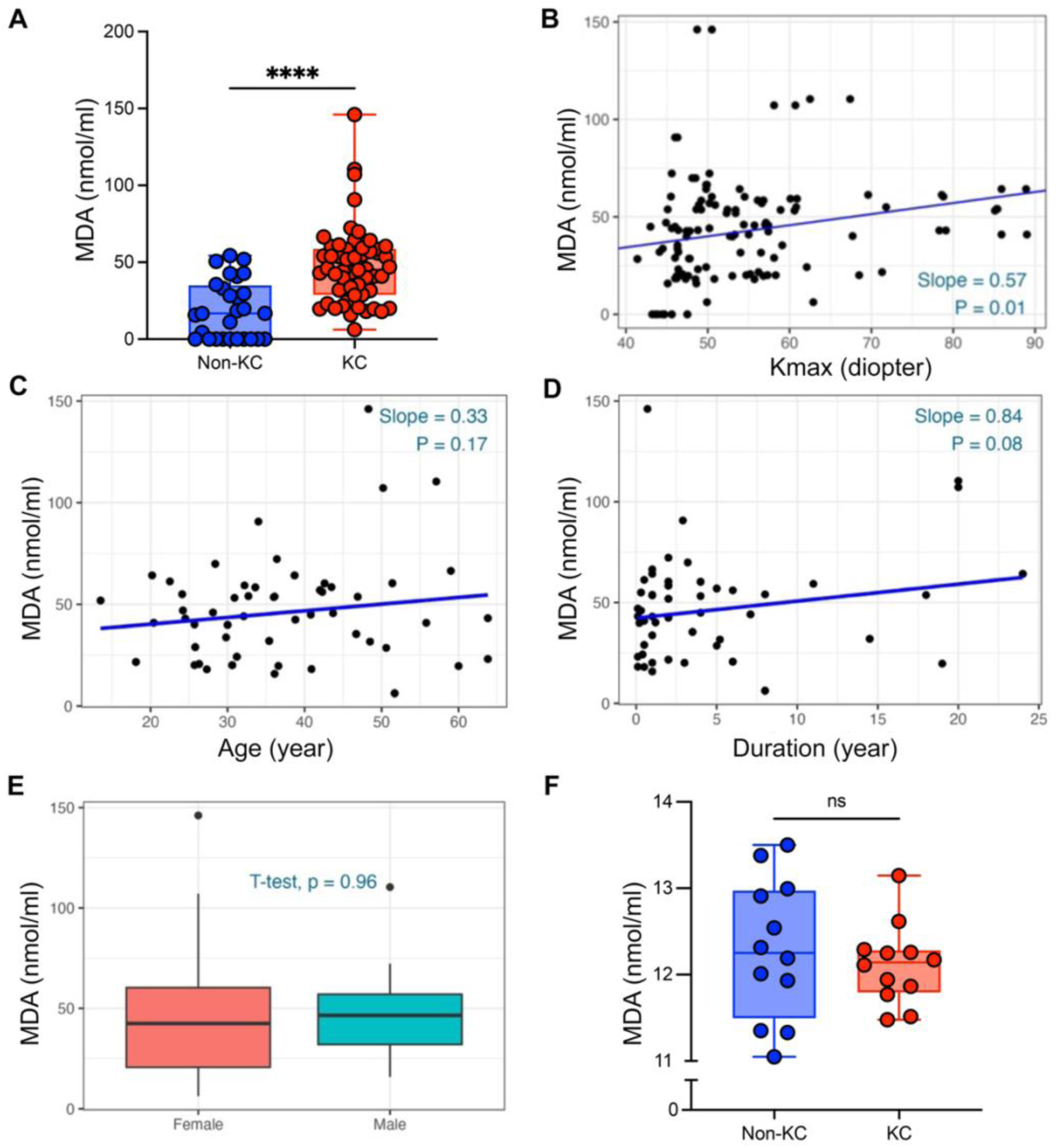
MDA elevated in KC patient tears. **(A)** MDA ELISA measurements on pooled tears per individual with KC (n = 53) or controls without KC (n = 28). Two-sample *t*-test between KC and non-KC groups. *p* < 0.001. **(B)** Linear regression analysis between subject-level tear fluid MDA levels and eye-level Kmax values in individual KC subjects show a positive correlation. **(C-D)** Tear MDA in KC subjects show no correlation with age or duration of disease. **(E)** No correlation between Tear fluid MDA and sex of participants. **(F)** Plasma MDA quantified by ELISA in KC (n = 12) and control non-KC (n = 12) subjects. A two-sample *t*-test shows no significant difference in plasma MDA levels between KC and non-KC. **(A, E & F)** Error bars = one SD. Two-sample *t*-test. **** p<0.0001, ns = non-significant.

### GPX3 in KC tear fluid correlates positively with Kmax and disease severity

We used 2.5 μg of total tear protein per sample **(Supplementary Table 4)** to determine GPX3 levels in 52 KC and 34 non-KC control samples (**Figure 2**). The GPX3 levels in KC and non-KC tear fluids were 2.57 ± 1.69 μg/ml and 0.92 ± 0.97 μg/ml, respectively, with the levels in KC being significantly higher than those of non-KC controls (p<0.001, two-sample type *t*-test (**Figure 2A**). Again, a linear regression analysis performed as described for MDA shows a strong positive correlation between GPX3 levels and Kmax values (**Figure 2B**). In KC patients stratified for disease severity (based on Kmax) we observed a significant difference between non-KC and moderate (p=0.01, ANOVA with Tukey post hoc tests), moderate and severe (p= 0.02) and non-KC and severe (p< 0.001) (**Supplementary Table 5**). We found no correlation between GPX3 levels and age (p=0.55; **Figure 2C**), disease-duration (p=0.62; **Figure 2D**), or sex (p=0.75; **Figure 2E**). GPX3 levels in plasma samples (**Figure 2F**) of a subset of individuals showed a slight increase (p=0.079) in KC (0.54 ± 0.307 μg/ml; n=11) compared to non-KC control ((0.34 ± 0.226 μg/ml; n=12).

**Figure 2:**
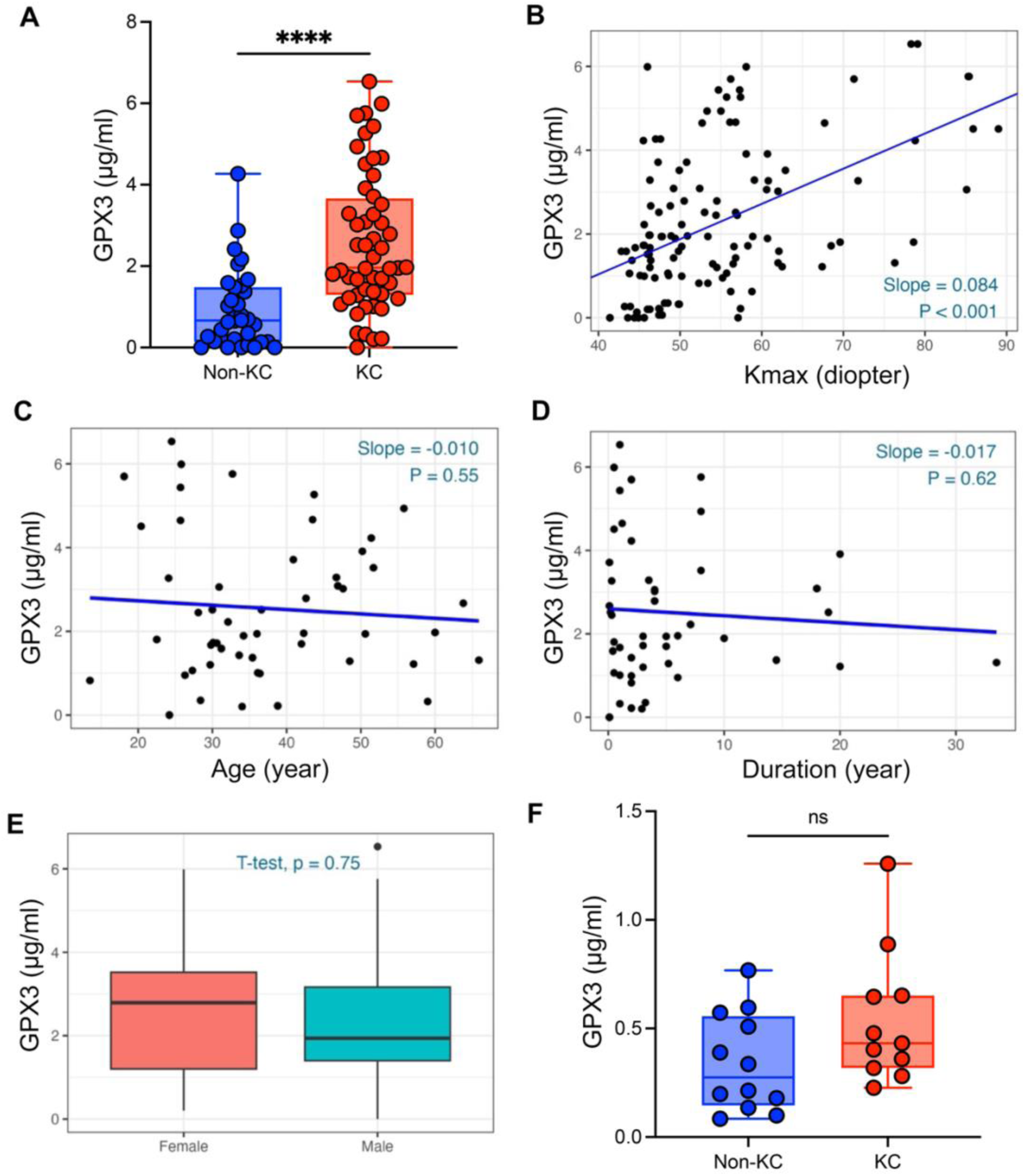
GPX3 elevated in KC patient tears and correlates positively with corneal steepness. **(A)** GPX3 protein measured by ELISA in pooled tear fluid per individual with KC (n = 54) and without KC (n = 32). **(B)** Linear regression analysis between subject-level tear fluid GPX3 and eye-level Kmax values per individual KC subject shows a positive correlation. **(C & D)** Linear regression analyses show no correlation between tear fluid GPX3 and age or duration of disease. **(E)** No sex-based differences in tear fluid GPX3 in KC patients. **(F)** Plasma GPX3 levels measured by ELISA in KC (n = 12) and non-KC (n = 12) individuals. **(A, E & F)** Error bars = one SD. Two-sample *t*-test. **** p<0.0001, ns = non-significant.

Since there are no reports of GPX3 in the cornea, we sought to investigate the presence of GPX3 in corneal tissue sections by immunohistochemistry. In donor corneas we detected strong immunostaining of GPX3 in the cytoplasm of all epithelial layers, and in a few stromal keratocytes in the stroma (**Figure 3A**). The donor cornea endothelial layer showed remarkably strong immunostaining for GPX3. Surgical KC corneal tissues did not have the endothelial layer, but we detected GPX3 immunostaining of the epithelial layers and some staining of anterior stromal keratocytes (**Figure 3B**) additional images in **Supplementary Figure 1**. Overall, our findings indicate that GPX3 is robustly produced by the corneal epithelial and endothelial layers. However, based on the limited number of donor and keratoconus samples we examined, it is difficult to ascertain any differences in GPX3 staining pattern between affected and unaffected corneas.

**Figure 3:**
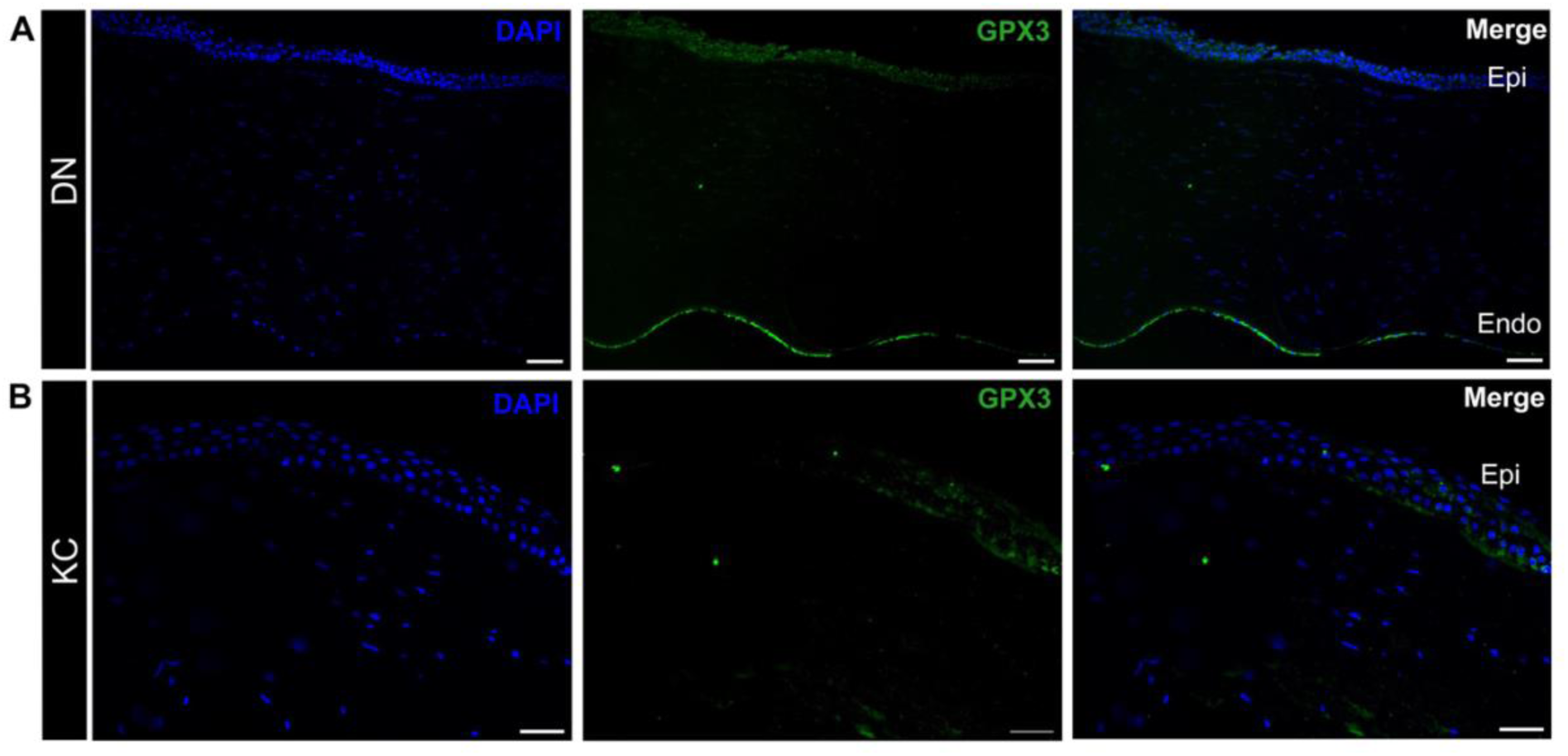
Immunolocalization of GPX3 in human corneal tissue. Representative immunofluorescence images showing GPX3 localization. **(A)** Immunostaining of GPX3 in DN corneas (n=2) localize GPX3 in epithelial layers and few stromal keratocytes, and the endothelium. **(B)** In KC corneas (n=2) GPX3 immunostaining was detectable in some epithelial cells and the basement membrane. The KC corneal samples did not have the endothelial layer. See Supplementary Figure 1 for additional DN and KC corneas. Scale bar = 20μm.

### Increased apoptotic cell death and suboptimal induction of antioxidants in keratoconus than in control donor stromal fibroblast cultures

We next asked whether response to H_2_O_2_-mediated oxidative stress would be impaired in cultured stromal fibroblasts from keratoconus compared to control donor cells. Second, we tested the effects of NRF2 signal-enhancing sulforaphane (SFN) in this cell culture model of oxidative stress (**Figure 4A**). Plated cells were pretreated with or without SFN and exposed to H_2_O_2_, followed by immunostaining for NRF2. Untreated DN and KC cells showed overall muted NRF2 staining of the entire cell body, but the staining was noticeably faint in KC cells (**Figure 4B**). With H_2_O_2_ treatment alone, we observed some nuclear staining of NRF2 in KC cells, and widespread, prominent nuclear staining of DN cells. The SFN-pretreatment and H_2_O_2_ combination showed increased nuclear staining of NRF2 in KC and DN cells. These results indicate that while KC fibroblasts can still activate NRF2 upon oxidative challenge, their reduced baseline NRF2 levels may increase susceptibility to damage. The ability of SFN to enhance NRF2 translocation and protect fibroblasts highlights the functional relevance of this pathway in maintaining stromal resilience.

**Figure 4:**
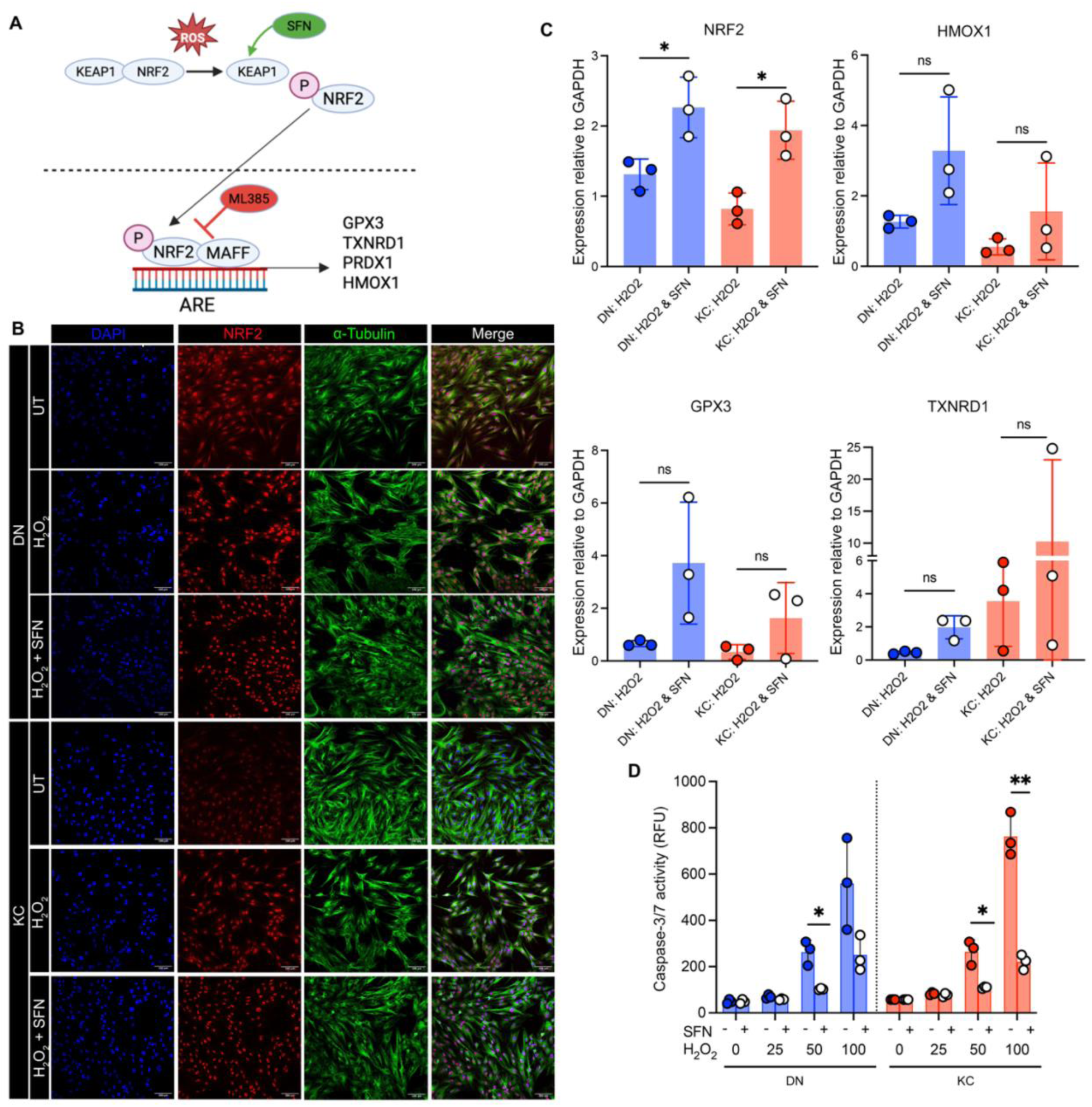
Impaired NRF2 nuclear translocation and enhanced apoptosis in KC stromal fibroblasts, rescued by Sulforaphane treatment. **(A)** A schematic showing NRF2 signal regulations. **(B)** Corneal fibroblasts derived from three biologically distinct DN and KC corneal tissues were treated with 10 μM SFN for 16 hours, followed by 50 μM H_2_O_2_ for 1 hour. Representative images of NRF2 immunostaining on untreated, H_2_O_2_-treated, and SFN + H_2_O_2_-treated cells. NRF2 shows stronger nuclear localization in H_2_O_2_-treated DN than in KC fibroblasts. H_2_O_2_ and SFN-pretreated cells shows strong NRF2-nuclear staining in DN and KC cells. Scale bar = 100 μm. Also see Supplementary Figure 3. **(C)** Gene expression of *NRF2* and its target genes, *HMOX1*, *GPX3* and *TXNRD1* in three distinct biological DN and KC fibroblasts treated with 10 μM SFN for 16 hours, followed by 50 μM H_2_O_2_ for 1 hour. Gene expression was determined by Taqman qRT-PCR and presented as ΔΔ cT normalized to GAPDH expression. *NRF2* expression was significantly upregulated in SFN + H_2_O_2_-treated cells compared to untreated cells but was not significantly different between DN and KC. **(D)** Caspase-3/7 activity was measured in DN and KC fibroblasts (n=3 biological replicates in each group) treated with 10 μM SFN for 16 hours, followed by increasing doses of H_2_O_2_. At the highest 100 μM H_2_O_2_ dosage the KC fibroblast stocks showed a significant increase in Caspase3/7 proapoptotic cells compared to DN. Ordinary one-way ANOVA (C) and two-way ANOVA test (D). *P < 0.05, **P < 0.001, ns = non-significant.

We used Taqman qRT-PCR to elucidate gene expression of *NRF2* and selected antioxidants*, GPX3*, *HMOX1* and *TXNRD1* in the SFN and H_2_O_2_-treated KC and DN cells (**Figure 4C**). The combined SFN and H_2_O_2_ treatment yielded a significant increase in GAPDH-normalized NRF2 expression in both DN and KC cells compared to H_2_O_2_ treatment alone, but there was no significant difference between DN and KC. Expression of the three antioxidants tested increased in SFN and H_2_O_2_-treated cells over H_2_O_2_ alone without reaching significance. These findings indicate that although KC fibroblasts show impaired induction of key antioxidant genes under oxidative stress, they remain responsive to pharmacological activation of NRF2 by SFN.

We sought to determine how cell survival and apoptosis would be affected by H_2_O_2_-mediated oxidative stress in KC and DN fibroblasts in this model (**Supplementary Figure 2** and **Figure 4D**). Plated adherent cells, with or without SFN pretreatment, were treated with increasing doses of H_2_O_2_ (0-100 𝜇M) and then assayed for pro-apoptotic caspase3/7 activity. H_2_O_2_ treatment alone increased Caspase 3/7 activity at all the H_2_O_2_ doses tested, with the highest dose causing a marked increase in pro-apoptotic cells in KC compared to DN fibroblasts. SFN pretreatment reduced Caspase-3/7 activity in KC and DN cells, suggesting similar rescue of apoptotic cells.

### NRF2 inhibition in DN fibroblast cultures evoke KC fibroblast-like phenotype

NRF2 appears to provide a central protective role in stromal cells. Therefore, we posited that inhibition of NRF2 in control DN fibroblasts would result in these cells to display keratoconus-like phenotype. ML385 is a benzoyl group-carrying small molecule that specifically binds NRF2 and interferes with the NRF2-MAFG complex from interacting with promoter regions of genes (29). We tested the effects of ML385 on the expression of selected antioxidant genes in six biologically distinct DN fibroblast stocks (**Supplementary Table 6**). We detected a decrease in the expression of all antioxidant genes tested, shown as fold change (FC), with the decrease in *HMOX1* (FC = 0.21 ± 0.12) and *PRDX1* (FC = 0.44 ± 0.32) being statistically significant (**Figure 5A**). We further tested the secretion of PRDX1 and GPX3 in the media and found these to decrease in ML385-treated versus DMSO vehicle-treated cells, where the decrease in PRDX1 was significant (p<0.008; **Figure 5B**). Morphologically, ML385-treated cells appeared more spindle-shaped, smaller and less spread out than the typical flattened fibroblastic morphology of untreated controls (**Figure 5C**) and showed significantly reduced growth in culture by seven days (**Figure 5D**).

**Figure 5:**
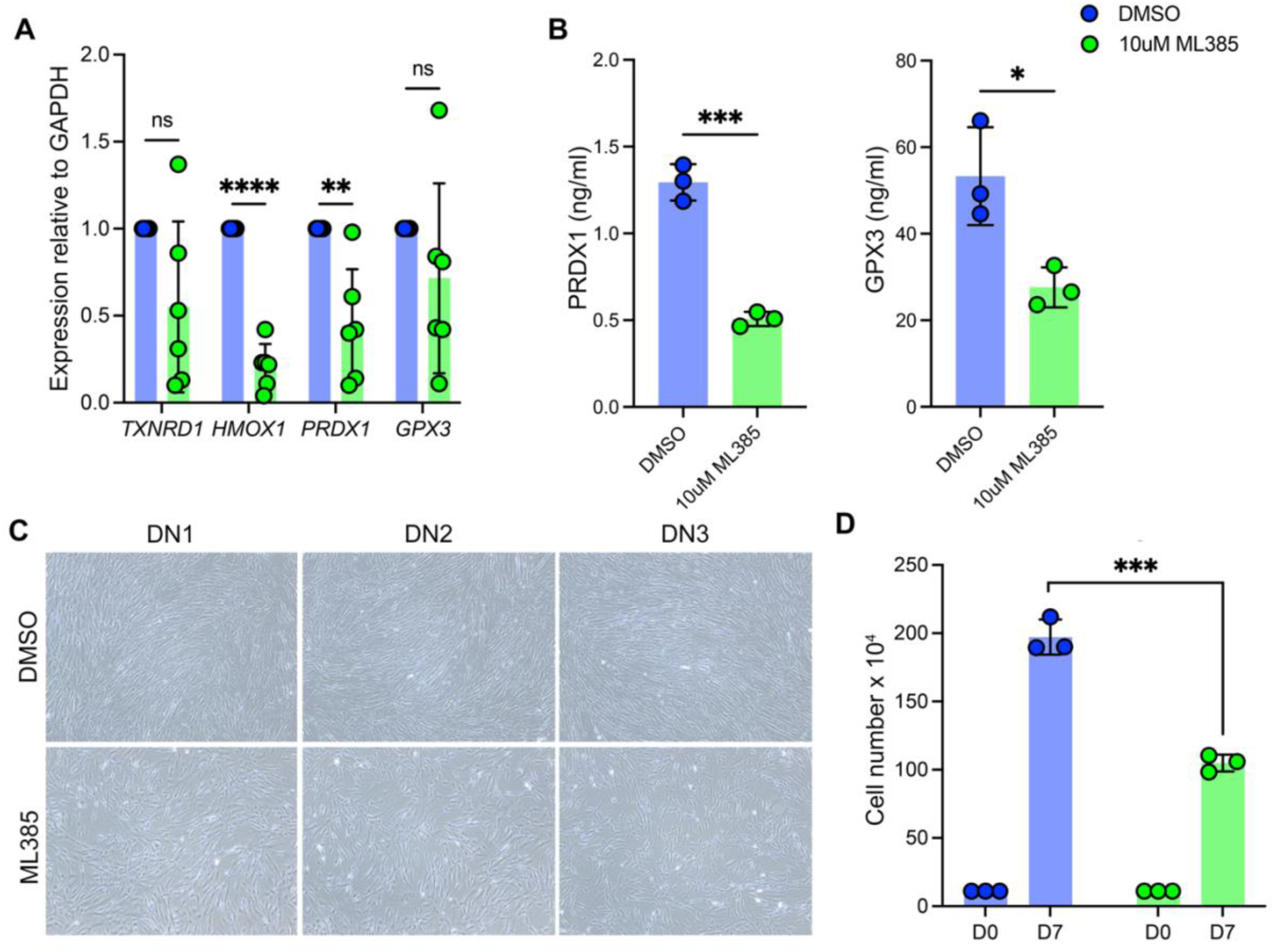
ML385-mediated NRF2 inhibition in DN fibroblasts suppresses antioxidant gene expression and cell growth. **(A)** Relative expression of *TXNRD1*, *HMOX1*, *PRDX1* and *GPX3* decrease in DN fibroblasts treated with 10 μM ML385 for 24 hours compared to DMSO (vehicle) controls. The results show mean ± SD from 6 biological replicates. **(B)** GPX3 and PRDX1 secreted into the culture media show a statistically significant decrease in DN fibroblasts treated with ML385. (n=3 biological replicates). Error bar shows mean ± SD. **(C)** Brightfield images of DN fibroblasts treated with ML385 or DMSO as a control. **(D)** Trypan blue-negative viable cells from **(C)** after seven days in culture. Student’s *t*-test, *p < 0.05, **P < 0.001, ***P < 0.001, ****P < 0.0001, ns = non-significant.

To assess whether this reduced growth was in part due to impaired proliferation, we tested BrdU incorporation in culture. Anti-BrdU-immunostaining (**Figure 6A**) revealed visibly fewer actively replicating BrdU-positive cells in the ML385-treated biological replicates compared to untreated controls. Quantitative assessments from 5 field of view show consistent decreases in 3 independent DN fibroblast stocks (**Figure 6B**). We also tested cell proliferation using Ki-67 immunostaining and detected a notable decrease in Ki-67–positive nuclei in ML385-treated fibroblasts (**Figure 6C**). These results indicate that inhibition of NRF2 inhibition compromises both antioxidant defense and fibroblast growth.

**Figure 6:**
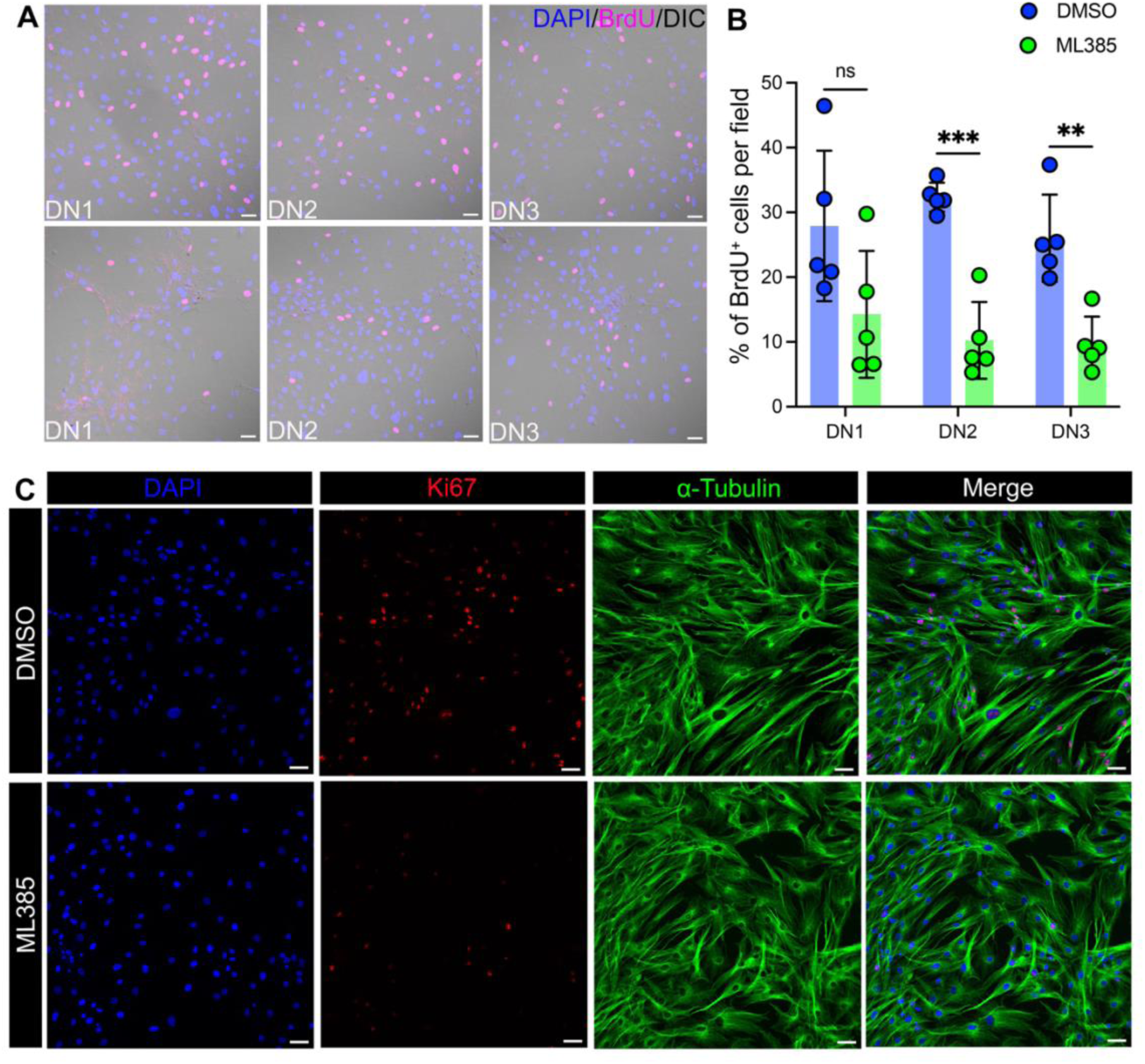
NRF2 inhibition in DN stromal cells evokes KC-like phenotype. **(A)** Representative immunofluorescence images of BrdU-positive cells, and **(B)** quantification of the percentage of BrdU-positive cells in DN fibroblasts (each dot represents an individual field). **(C)** Immunofluorescence images showing Ki-67 staining in DN fibroblasts (Representative images from one of three biological replicates) Data are presented as mean ± SD. Statistical analysis was performed using Student’s *t*-test, **P < 0.01, ***P < 0.001, ns = non-significant. Scale bar = 50μm.

Corneal keratocyte and fibroblast health is intricately connected to ECM production, including collagen type I, V, proteoglycans and tissue remodeling matrix metalloproteinases (30–32). Therefore, we examined whether NRF2 inhibition would perturb ECM deposition. We tested the deposition of fibrillar collagen types I and V and fibronectin in ML385-treated DN fibroblasts. We observed a remarkable decrease in COL1A1 and COL5A1 immunostaining after ML385-mediated inhibition of NRF2 (**Figure 7A & B**). Immunostaining of fibronectin was slightly reduced in the ML385-treated DN fibroblasts (**Figure 7C**). Expression of *COL1A1* and *COL5A1* determined by qPCR indicated a decrease in both, (FC = 0.29 ± 0.28) and (FC = 0.61 ± 0.23), respectively, following NRF2 inhibition (**Figure 7D**). Together, these results show that NRF2 is essential for maintaining ECM gene expression and matrix organization in corneal fibroblasts. Further, ML385-treated DN fibroblasts may likely be useful as a keratoconus cell culture disease model.

**Figure 7:**
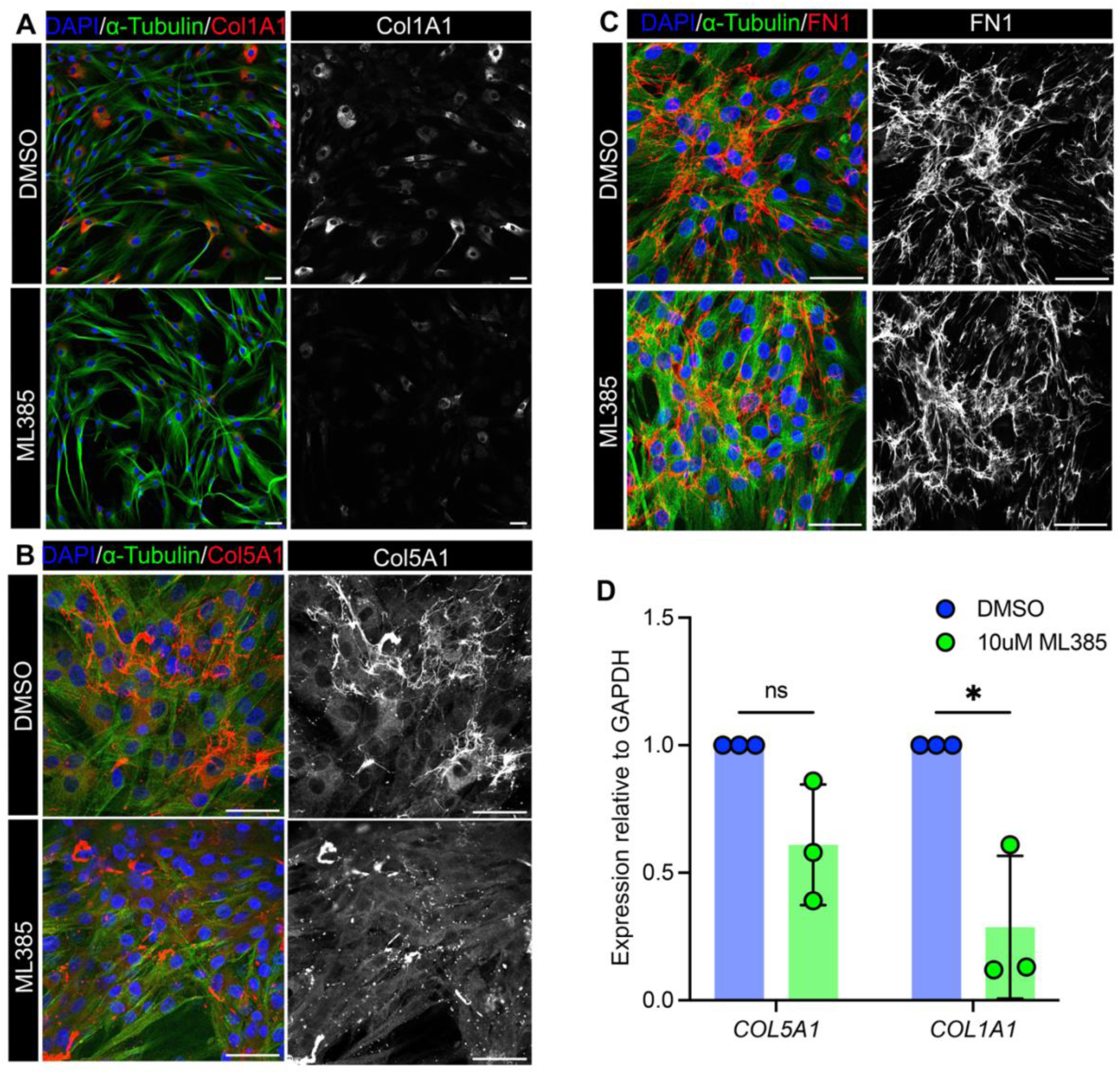
NRF2 inhibition causes reduced ECM in DN fibroblasts. **(A-C)** Immunofluorescence staining images showing the corneal fibroblasts from three distinct donors were treated with DMSO (vehicle control) or ML385 for COL1A1 **(A)**, COL5A1 **(B)**, and FN1 (Fibronectin-1) **(C)**. Representative images from one of three biological replicates. **(D)** Bar graphs representing the gene expression analysis of *COL1A1* and *COL5A1* in DMSO- and ML385-treated cells. Data are presented as mean ± SD, (n=3 biological replicates). Statistical analysis was performed using Student’s *t*-test, *P < 0.05, ns = non-significant. Scale bar = 50μm.

## Discussion

We found consistent elevations in MDA levels in the tear fluid of keratoconus patients collected by the capillary tube method. Using a different Schirmer strip method for sampling tear fluids, another study reported increased MDA in a smaller set of KC patients (33). In a subset of subjects for whom we have tear fluid data, we measured MDA in the plasma and found no difference between KC and non-KC controls. Thus, the MDA accumulation appears to occur only in the ocular environment. A recent study investigated correlations between circulating plasma MDA levels in a case-control study of KC patients from Iran with a 50 base-pair deletion polymorphism in the promoter region of SOD1 (34) that reduces SOD1 transcript and reportedly also correlates with cardiovascular disease (35). The SOD1-KC study found significant increases in circulating MDA in KC patients heterozygous or homozygous for the SOD1 promoter polymorphism. While we have not genotyped our cohorts for the SOD1 polymorphism, our KC patients are mostly of European and African ancestry, and unlikely carriers of this polymorphism.

We detected a consistent and statistically significant increase in GPX3 levels in the tear fluid of KC patients. High Kmax measurements indicate steep corneal curvature and increased keratoconus disease severity. Therefore, a strong positive correlation of tear fluid GPX3 levels with Kmax suggests a correlation with disease severity. However, only a small number of controls visiting the eye clinic underwent pachymetry imaging of the ocular surface. Therefore, we were unable to test its correlation with corneal steepness in keratoconus-unaffected subjects. Assaying GPX *activity*, but not GPX3 levels specifically, in a limited number of KC patients before/after UV crosslinking treatment, the Balmus group reported an increase in total GPX activity in KC patients after UV crosslinking (33). Our study indicates a strong correlation between KC, MDA and GPX3 protein levels, where the latter, may become useful as a tear fluid biomarker for KC. However, the underlying mechanisms that regulate GPX3 levels in the tear fluid remain poorly understood. The GPXs are selenoproteins, largely investigated in the context of cancer biology (36). Of the 8 GPXs, GPX3 is secreted, and remains attached to the basement membrane of tissues (37), which fits well with our specific immunostaining of GPX3 at the corneal epithelial, endothelial surfaces and the pericellular matrix of epithelial, endothelial and keratocyte cells. The GPX3 promoter carries a metal recognition element (MRE) and an antioxidant response element (ARE) recognized by NRF2 and is also regulated by the hypoxia-inducible factor (HIF-1). Thus, even though our studies suggested reduced NRF2 activity in KC corneas, compensatory increases in other regulatory signals can explain the GPX3 elevations we see in KC tear samples. The bulk of proteins in the aqueous layer of the tear fluid originate from the lacrimal gland. Therefore, ocular surface perturbations in KC, such as allergy, eye rubbing and dry eyes, can lead to altered tear fluid levels and composition in keratoconus, and this may also underlie the positive correlation we see with KC disease severity and tear fluid GPX3. We further recognize that, the control group in our tear analysis included individuals with ocular conditions such as glaucoma suspicion, cataract, and dry eye. However, due to the inherent challenges in recruiting completely ocular-disease–free individuals, inclusion of subjects with mild ocular comorbidities was an unavoidable aspect of clinical sample recruitment. To bridge this limitation, in vitro experiments were conducted to validate NRF2-mediated antioxidant responses under controlled experimental oxidative stress.

The tear fluid is accessible with relative ease and provides an insight into the ocular environment. While the total yield in volume is quite limited, it is modestly rich in proteins, ranging from 6-10 mg/ml based on early proteomic studies (38). We and others reported increases and decreases in selected proinflammatory cytokines in the tear fluid of keratoconus patients (39, 40). Prolactin-inducible protein (PIP), another potential biomarker was detected as downregulated in keratoconus (41). Interestingly, PIP showed concordant decreases in plasma and urine of keratoconus patients and is reportedly regulated by sex hormones (42). Our MDA and GPX3 findings are different from PIP in that they are not similarly regulated in plasma and show no sex-based differences.

The ultimate thinning of the cornea and decreased stromal cell counts in KC prompted us and many others to examine stromal ECM and keratocyte health in freshly isolated cells from KC corneas (43–45). In the current study, we specifically show that KC stromal cell cultures undergo increased Caspase-3/7 mediated apoptosis when challenged with H_2_O_2_-mediated oxidative stress, while treatment with SFN restores cell survival in KC cells to DN-levels. Previously, we found that stromal keratocytes isolated from KC corneas exhibited enhanced non-canonical TGF-β signaling (43). We also found that KC corneal stromal cells cultured in low glucose serum-free media produced lower amounts of assembled ECM collagens, but increased activation of MMP2 and expression of integrated stress response markers (45). Overall, these studies show loss of homeostasis at basal states in KC and compared to DN cells, and the former were less successful in overcoming experimental oxidative stress. Finally, we tested if DN stromal keratocytes could acquire KC-like diseased phenotype upon blocking NRF2 activation with ML385. DN cells treated with ML385 showed poor activation of NRF2, decreased secretion of antioxidants, reduced cell growth and lower fibrillar collagen deposition. While our cell culture studies used 5-6 biologically independent KC and DN cell cultures, additional testing of corneal keratocyte cell cultures from distant groups would increase scientific rigor.

Corneal endothelial degeneration in Fuchs dystrophy has also been linked to unresolved oxidative stress (27, 46). Further, the Jurkunas team showed that NRF2 activity and endothelial cellular apoptosis in *ex vivo* models of oxidative stress could be rescued with sulforaphane treatment (46, 47). Our cell culture models also suggest that antioxidant functions in keratocytes could be improved with sulforaphane, a potential therapeutic supplement for keratoconus patients.

Our finding shows that both MDA and GPX3 levels increase in the tear fluid make a compelling case for developing these as biomarkers that may help to detect keratoconus before overt corneal ectasia becomes evident. Further studies could also lead to the use of these oxidative stress-related factors to track disease progression or response to treatments. A limitation of this study is that our patients were all from the same geographical area. Additional testing of cohorts from other parts of the world is required to determine whether these findings are universal.

## Methods

### Study approval

This study was approved by the Institutional Review Board of NYU Grossman School of Medicine and conducted in accordance with the Declaration of Helsinki. All subjects provided written informed consent. Keratoconus patients filled out a questionnaire regarding disease duration, other eye conditions such as allergy and atopy (**Supplementary Figure 4**).

### Sex as a biological variable

This study examined male and female individuals and the distribution of males and females in the study groups were not markedly different. In the control participants there were 15 males and 19 females. The keratoconus patient group contained 25 males and 28 females (**Supplementary Table 1**). Differences in tear fluid MDA and GPX3 levels between male and female participants were assessed to evaluate sex-specific variations.

### Study participants

57 individuals with keratoconus (KC) and 34 without keratoconus (non-KC) were included (**Supplementary Table 1**). Unaffected control volunteers were those visiting the NYU Langone Eye Center for routine eye examination, glaucoma and myopia assessments. KC severity was assigned based on keratometry (𝑘) measurements, with mild as steepest 𝑘 <45 diopter (D), moderate 45D ≥steepest 𝑘 ≤ 52D and severe as steepest 𝑘 > 52.

### Tear fluid and peripheral blood plasma

Tear samples, 5-8 𝜇𝑙 per eye were collected using micro glass capillary tubes (P0674, Millipore sigma) from the cul-de-sac of the eye without inducing reflex tear as in our previous study (40). Tear fluid from both eyes were added to PBS, centrifuged (1000 rpm 5 minutes) to remove the cell debris and stored at −80°C until further use. Total protein was measured using the Nanodrop One (Thermo Scientific). Peripheral blood collected by standard phlebotomy techniques were used to extract plasma and stored at −80°C till use.

### ELISA

GPX3 (IT3662, G-Biosciences) and MDA (STA-832, Fischer Scientific) were measured using ELISA in tear and plasma samples following manufacturer’s recommendations. Tear samples were diluted such that 2.5μg (for GPX3) and 5μg (for MDA) of total protein were loaded into each well. Final concentrations of MDA and GPX3 were calculated from the standard curve for each kit.

### qRT-PCR

Total RNA was extracted using TRIzol reagent (Thermo Fisher Scientific) and reverse transcribed using iScript cDNA synthesis kit, and quantitative PCR (qPCR) was performed using Fast one step Taqman master mix (BIO-RAD) on a QuantStudio 5 Real-Time PCR System for genes listed in **Supplementary Table 7**. The following thermal cycling conditions were used, initial denaturation at 95°C for 2 minutes, followed by 40 cycles of 95°C for 15 seconds, 60°C for 30 seconds, and 72°C for 30 seconds. GAPDH-Relative expression was presented as ΔΔCt.

### Corneal stromal cell extraction and culture

Stromal cells were extracted from donor corneas and discarded KC corneal tissue (**Supplementary Table 6**) as in our previous studies (43, 48). Briefly, the corneas were washed with PBS, the epithelial and endothelial layer (in case of donor corneas) were removed mechanically, the remaining stroma was minced into pieces and transferred to 3.3 mg/ml collagenase type I (#17100017, Invitrogen) in DMEM media and incubated at 37°C with shaking (125 rpm). Breakdown of the tissue was tracked visually every 30-60 minutes, when the corneal pieces were allowed to settle under gravity and the supernatant transferred to a fresh tube on ice, adding fresh collagenase solution to the indigested material. Most of the tissue was digested within 2-4 hours. The cell suspension was centrifuged at 1500 rpm for 5 minutes, and the cell pellet was resuspended in DMEM F12, 10% FBS, 1X Antibiotic and antimycotic, and used for up to 7 passages.

### In vitro oxidative stress

DN and KC stromal fibroblasts were cultured in DMEM/F12 supplemented with 10% FBS and 1% penicillin-streptomycin at 37°C with 5% CO_2_. We used DN fibroblasts to test cell viability in 0-100 µM H_2_O_2_ in the culture media for one hour and selected 50 µM H_2_O_2_ as the optimal dose. For the oxidative stress experiments, DN and KC fibroblast stocks were pre-treated with or without 10 µM sulforaphane for 16 hours, followed by 50 µM H_2_O_2_ for 1 hour. To inhibit NRF2 binding to AREs of target genes, cells were treated for 24 hours with 10 µM final concentration of ML385 from DMSO dissolved stock solutions to inhibit NRF2 binding to ARE and target gene activation.

### Cell viability assay

Cell viability was assayed using the resazurin reduction as we described previously (45). Briefly, DN and KC fibroblasts in a 96-well plate were incubated with alamar blue/resazurin (#R7017; Sigma-Aldrich) for 1 hour at 37°C, and fluorescence intensity was measured using a microplate reader (excitation: 530–560 nm; emission: 590 nm). The fluorescence intensity was directly proportional to the metabolic activity of viable cells. Cell numbers in the test wells were estimated using a standard curve we generated with known cell numbers.

### Immunofluorescence on human corneal fibroblasts

The fibroblasts cultured in chamber slides were fixed with 4% paraformaldehyde for 15 minutes and permeabilized using 0.5% Triton X-100 in PBS for 10 minutes. Slides were blocked with 5% bovine serum albumin (BSA) for 1 hour and incubated overnight at 4°C with primary antibodies (**Supplementary Table 7**). After washing, fluorophore-conjugated secondary antibodies were added for 1 hour at room temperature in the dark. Nuclei were stained with DAPI, and slides were mounted using anti-fade medium. Images were acquired with a confocal microscope (LSM 880, Zeiss).

### Immunofluorescence on tissue sections

5 µm thick paraffin embedded tissue sections were deparaffinized, rehydrated, and subjected to antigen retrieval in citrate buffer (pH 6.0) at 95°C for 20 minutes. After blocking with 5% BSA, the sections were incubated overnight at 4°C with primary antibody (**Supplementary Table 7**). After incubating with HRP-conjugated secondary antibodies, the slides were counterstained with hematoxylin, dehydrated, and mounted, and images acquired in a confocal microscope (LSM 880, Zeiss).

### Assessment of Caspase-3/7 activity

Apoptotic cells were quantified using the Apo-ONE Homogeneous Caspase-3/7 Assay Kit (Promega, USA) as we described previously (49). In brief, 10,000 fibroblasts (DN=3; KC=3) were seeded per well in 96-well plates. After overnight attachment, cells were treated with H_2_O_2_-induced oxidative stress followed by the addition of the Apo-ONE Caspase-3/7 reagent. The plates were incubated at 37°C for 18 hours with constant agitation at 300 rpm to enable substrate cleavage. Fluorescence intensity was recorded at excitation/emission wavelengths of 485/530 nm using a fluorescence microplate reader (FlexStation® 3 Reader), and relative fluorescence units (RFU) were analyzed with SOFTmax Pro software (Molecular Devices).

### BrdU assay

Proliferating cells were detected using the 5-bromo-2’-deoxyuridine (BrdU) incorporation approach. In brief, cells were incubated with 0.03 mg/mL BrdU in pre-warmed growth medium for 24 hours at 37°C. Cells were fixed with cold 70% ethanol for 5 minutes at room temperature, washed with PBS, incubated in 1.5 M HCl for 30 minutes followed by additional PBS washes, incubated in blocking buffer followed by overnight incubation with anti-BrdU primary antibody at 4°C, anti – mouse 488 secondary antibody for 45 minutes, washed with PBS washes, and counterstained with DAPI and visualized by fluorescent microscopy. The percentage of BrdU positive cells was calculated in reference to DAPI-stained nuclei using ImageJ.

### Statistical analyses

Since GPX3 and MDA were measured in pooled tear fluid from both eyes per person, these are subject-level variables. Therefore, a two-sample *t*-test was used to compare GPX3 or MDA between individuals with or without keratoconus. We also used One-way ANOVA with Tukey post hoc tests to compare GPX3 or MDA levels across 4 groups: Non-KC, Mild-, Moderate-, and Severe-KC. To assess correlations between Kmax (eye-level variable) and GPX3 or MDA (subject-level variable), we conducted a linear regression analysis, including both eyes and using cluster-robust standard errors to account for within-patient correlation. This allowed use of all available eye-level data while providing valid p-value. For the *in vitro* experiments, Student’s *t* test was used to compare treatment and control groups with statistical significance set at p≤ 0.05.

### Data availability

All data will be made available in the main and supplementary data files at the time of publication.

## Author Contributions

SC conceived the study, and SC, MAK, GM, and VS interpreted the data. SC and MAK wrote the manuscript. MAK performed tear fluid measurements and cell culture studies. GM assisted with analysis and figure preparations. MC managed sample handling and data management. Patient samples were provided by RM, LS, EHK, CRP, IDH, and ALB. Tear fluid statistical analyses were conducted by TFL. Immunohistochemistry on some of the donor and keratoconus corneas was performed by RS, RD, and VS.

## Acknowledgements

The authors are grateful to patient and control subjects for providing tear fluid samples. We thank the study coordinators, Jamika Singleton-Garvin, Sugeidy Ferreira Brito, Evan Shapiro, and Dzhalil Abizgildin for their assistance in the IRB protocols, consenting subjects, and collecting blood and tear fluid. We are thankful to the NYU Langone Microscopy Laboratory (RRID: SCR_017934), the Experimental Pathology Research Laboratory (RRID: SCR_017928), all of which receive partial support from the Laura and Isaac Perlmutter Cancer Center Support Grant P30CA016087 and the Shared Instrument Grant S10 OD021747. The study was funded by an NIH grant, R01EY026104 to SC. The authors thank Research to Prevent Blindness (RPB) for supporting the Department of Ophthalmology with an unrestricted grant.

## Notes

### Competing Interest Statement

The authors have declared no competing interest.

