## Supplemental information for "Oxidative-stress related increase in keratoconus tear MDA and GPX3 while NRF2-antioxidant functions decrease in stromal cells"

This file includes

Supplementary Table 4 to 7
Supplementary Figure 1 to 4

**List of Supplementary Tables**

**Supplementary Table 1:** Sample information of the tear samples used for measuring MDA and GPX3 (Data provided in excel sheet).

**Supplementary Table 2:** Tear protein concentrations for the samples used for measuring MDA (Data provided in excel sheet).

**Supplementary Table 3:** Statistical evaluation of tear MDA concentrations among keratoconus severity grades.

**Supplementary Table 4:** Tear protein concentrations for the samples used for measuring GPX3 (Data provided in excel sheet).

**Supplementary Table 5:** Statistical analysis of tear GPX3 concentrations across keratoconus severity groups.

**Supplementary Table 6**: Demographic characteristics and experimental utilization of study samples.

**Supplementary Table 7:** Reagent information.

**List of Supplementary Figures**

**Supplementary Figure. 1:** Immunolocalization of GPX3 in additional human corneal samples.

**Supplementary Figure. 2:** Dose optimization of $H_2O_2$ for inducing oxidative stress in corneal fibroblasts.

**Supplementary Figure. 3:** Higher-magnification view of NRF2 immunolocalization in corneal fibroblasts.

**Supplementary Figure 4**: Keratoconus study questionnaire.

**Supplementary Table 3:** Statistical evaluation of tear MDA concentrations among keratoconus severity grades.

**ANOVA with Tukey post hoc tests**

One-way ANOVA compared MDA levels across four groups (No KC, Mild, Moderate, Severe), followed by Tukey's post hoc tests.

| MDA | | | | p value (ANOVA) |
|---|---|---|---|---|
| Severity | n | mean | sd | |
| No | 29 | 20.16 | 19.10 | <0.001 |
| Mild | 8 | 39.10 | 15.33 | |
| Moderate | 31 | 47.79 | 29.65 | |
| Severe | 32 | 45.28 | 19.35 | |

**Tukey post hoc:** Significant differences were observed between No vs Moderate (p = 0.01), and Severe vs. No (p < 0.001). No significant differences were found between Severe vs. Mild, Moderate vs. Mild, Severe vs Moderate, or No vs. Mild.

| **Severity group** | **p adj** |
|---|---|
| Moderate-Mild | 0.77 |
| No-Mild | 0.17 |
| Severe-Mild | 0.90 |
| No-Moderate | <0.001 |
| Severe-Moderate | 0.97 |
| Severe-No | <0.001 |

**Supplementary Table 5:** Statistical analysis of tear GPX3 concentrations across keratoconus severity groups.

**ANOVA with Tukey post hoc tests**

One-way ANOVA compared GPX3 levels across four groups (No KC, Mild, Moderate, Severe), followed by Tukey's post hoc tests.

| GPX3 | | | | p value (ANOVA) |
|---|---|---|---|---|
| **Severity** | **n** | **mean** | **sd** | |
| **No** | 34 | 0.92 | 0.97 | <0.001 |
| **Mild** | 6 | 1.83 | 1.37 | |
| **Moderate** | 29 | 2.06 | 1.44 | |
| **Severe** | 33 | 3.11 | 1.78 | |

**Tukey post hoc:** Significant differences were observed between No vs Moderate (p = 0.01), Severe vs Moderate (p = 0.02), and Severe vs. No (p < 0.001). No significant differences were found between Severe vs. Mild, Moderate vs. Mild, or No vs. Mild.

| Severity group | p adj |
|---|---|
| **Moderate-Mild** | 0.98 |
| **No-Mild** | 0.48 |
| **Severe-Mild** | 0.19 |
| **No-Moderate** | 0.01 |
| **Severe-Moderate** | 0.02 |
| **Severe-No** | <0.001 |

**Supplementary Table 6**: Demographic characteristics and experimental utilization of study samples.

| Sample | Sex | Age | Ancestry | Experimental use |
|---|---|---|---|---|
| DN10605 | Female | 55 | Caucasian | NRF2 activation, NRF2 inhibition |
| DN21404 | Female | 63 | Caucasian | NRF2 inhibition |
| DN13842 | Female | 6 | Caucasian | NRF2 activation, NRF2 inhibition, |
| DN21568 | Female | 59 | Caucasian | NRF2 inhibition |
| DN15091 | Male | 63 | Caucasian | NRF2 activation, NRF2 inhibition |
| DN12091 | Un | Un | Un | NRF2 inhibition |
| DN921 | Un | Un | Un | NRF2 inhibition |
| DN1 | Male | 35 | Asian | GPX3 - IF |
| DN2 | Un | Un | Un | GPX3 - IF |
| DN3 | Female | 63 | Caucasian | GPX3 - IF |
| KC826 | Female | 29 | Black | NRF2 activation |
| KC2807 | Male | 76 | Caucasian | NRF2 activation |
| KC304 | Female | 23 | American Asian | NRF2 activation |
| KC1 | Female | 26 | Asian | GPX3 - IF |
| KC2 | Female | 38 | Asian | GPX3 - IF |
| KC3 | Un | Un | Un | GPX3 - IF |
| Un = Unknown, NRF2 = Nuclear factor erythroid 2-related factor 2, GPX3 = Glutathione peroxidase, IF = Immunofluorescence | | | | |

**Supplementary Table 7: Reagent information**

| Reagent/Kit | Source | Catalog Number | Application/Use |
|---|---|---|---|
| Micro glass capillary tubes | Millipore Sigma | P0674 | Tear sample collection |
| GPX3 ELISA kit | G-Biosciences | IT3662 | ELISA for tear and plasma samples |
| MDA ELISA kit | Fischer Scientific | STA-832 | ELISA for tear and plasma samples |
| TRIzol reagent | Thermo Fisher Scientific | 15596018 | RNA extraction |
| iScript cDNA synthesis kit | Bio-Rad | 1708891 | Reverse transcription |
| Fast One Step TaqMan master mix | Bio-Rad | 4444557 | Quantitative PCR |
| TaqMan gene assay: NFe2l2 | Thermo Fisher Scientific | Hs00975961_g1 | qRT-PCR |
| TaqMan gene assay: HMOX1 | Thermo Fisher Scientific | Hs01110250_m1 | qRT-PCR |
| TaqMan gene assay: TXNRD1 | Thermo Fisher Scientific | Hs00917067_m1 | qRT-PCR |
| TaqMan gene assay: GPX3 | Thermo Fisher Scientific | Hs01078668_m1 | qRT-PCR |
| TaqMan gene assay: COL1A1 | Thermo Fisher Scientific | Hs00164004_m1 | qRT-PCR |
| TaqMan gene assay: COL5A1 | Thermo Fisher Scientific | Hs00609133_m1 | qRT-PCR |
| TaqMan housekeeping gene: GAPDH | Thermo Fisher Scientific | Hs02758991_g1 | qRT-PCR normalization |
| Collagenase type I | Invitrogen | 17100017 | Corneal stromal digestion |
| DMEM/F12 medium | Invitrogen | 11330032 | Cell culture |
| Fetal bovine serum (FBS) | Gibco | 16140071 | Cell culture |
| Antibiotic/antimycotic solution | Thermo Fisher Scientific | 15240062 | Cell culture |
| Sulforaphane (SFN) | Med Chem Express | S6317 | NRF2 activation |
| Hydrogen peroxide ($H_2O_2$) | Millipore Sigma | H1009 | Oxidative stress induction |
| ML385 | Med Chem Express | HY-100523 | NRF2 inhibition |
| Alamar Blue / Resazurin | Sigma-Aldrich | R7017 | Cell viability assay |

| | | | |
|---|---|---|---|
| Paraformaldehyde 4% | Thermo Fisher Scientific | 28908 | Cell fixation |
| Triton X-100 0.5% | T-8787 | Sigma | Cell permeabilization |
| Bovine serum albumin (BSA) 5% | Fisher BioReagents | 9048-46-8 | Blocking |
| DAPI | BD Pharmingen | 564907 | Nuclear staining |
| Apo-ONE Caspase-3/7 Assay Kit | Promega | G7792 | Apoptosis measurement |
| BrdU 5-bromo-2'-deoxyuridine | Med Chem Express | HY-15910 | Cell proliferation assay |
| Anti-BrdU primary antibody | Cell Signalling Technology | 5292T | BrdU assay |
| Anti-mouse 488 secondary antibody | abcam | ab150117 | BrdU assay detection |

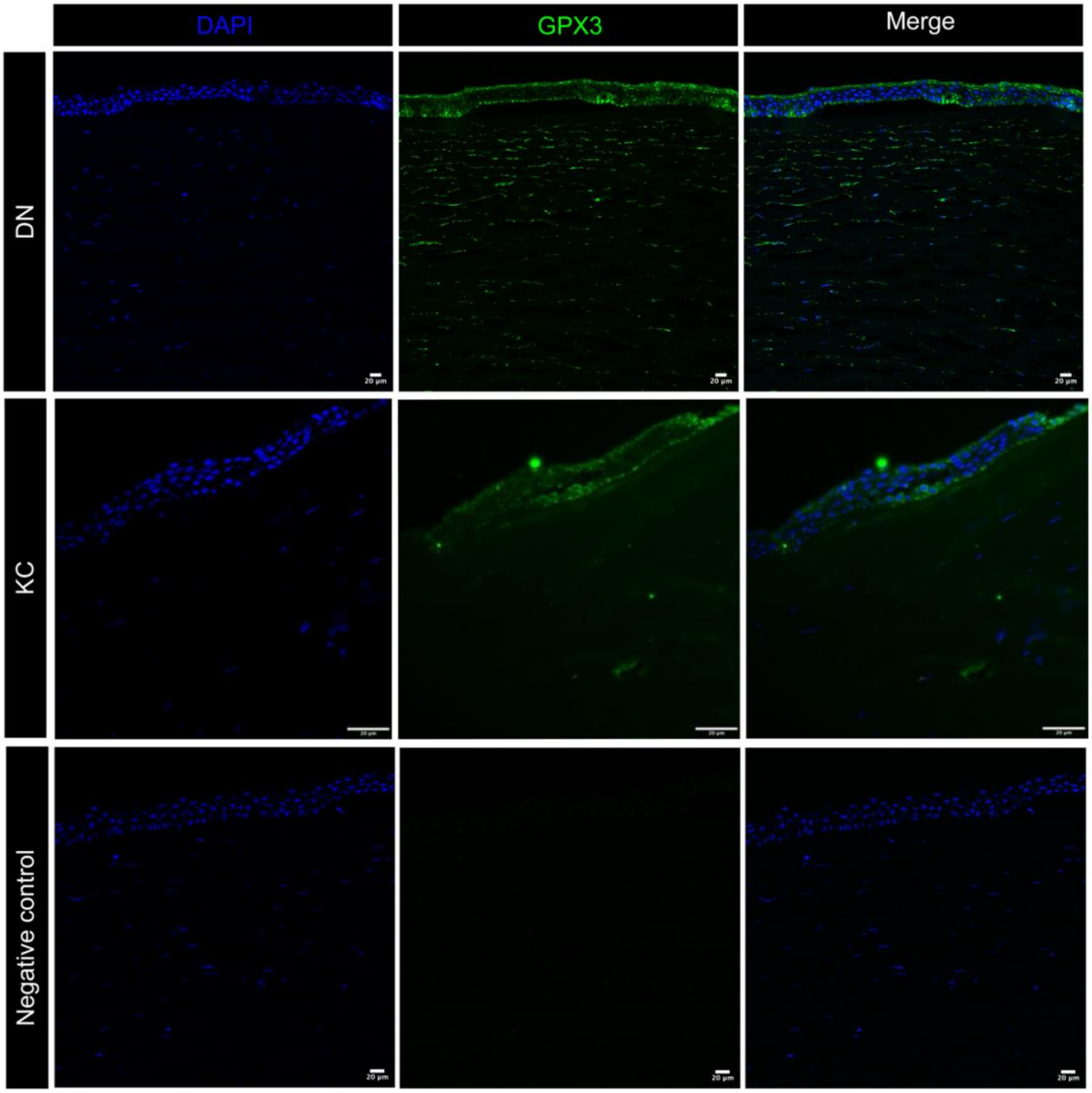


**Supplementary Figure. 1: Immunolocalization of GPX3 in additional human corneal samples.** Representative immunofluorescence images showing GPX3 expression in another set of donor corneas (DN2) and keratoconus corneas (KC2). Similar to the main figure, GPX3 was localized in the epithelial layers. (C) No-primary antibody control shows absence of non-specific staining, confirming the specificity of GPX3 immunolabeling.

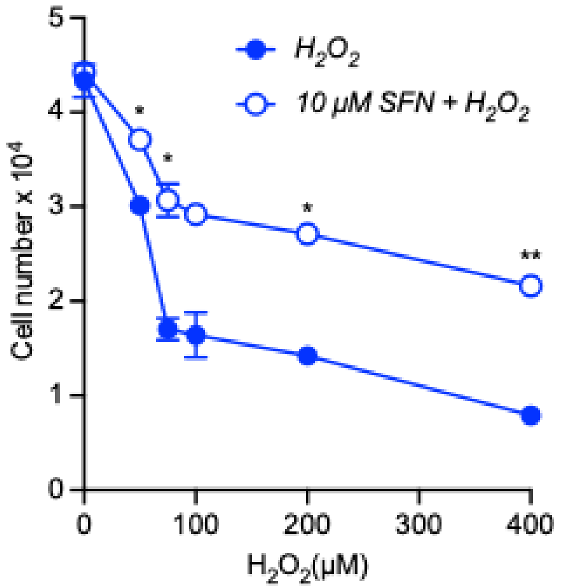


c

**Supplementary Figure. 2: Dose optimization of $H_2O_2$ for inducing oxidative stress in corneal fibroblasts.** Cell viability was assessed using the resazurin reduction assay following treatment with increasing concentrations of $H_2O_2$, with or without sulforaphane (SFN). Error bars represent [mean ± SD]. Statistical analysis was performed using the two-way ANOVA test. *P < 0.05, **P < 0.01.

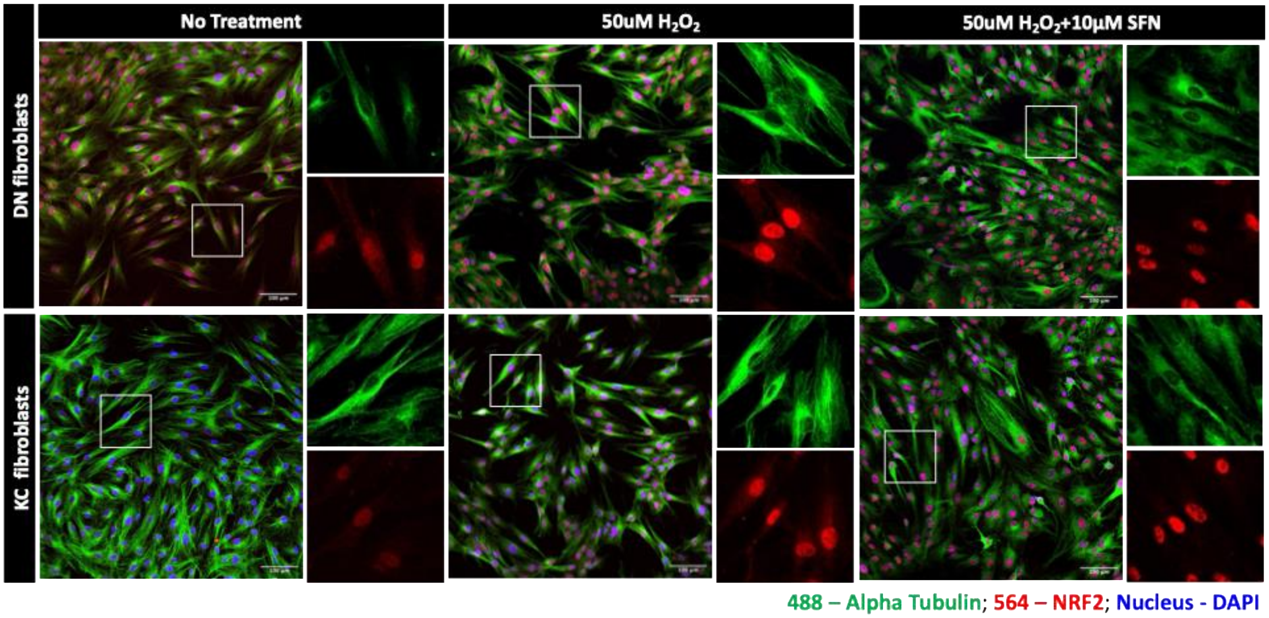


**Supplementary Figure. 3: Higher-magnification view of NRF2 immunolocalization in corneal fibroblasts.** Representative enlarged images corresponding to Figure 4, showing detailed visualization of NRF2 distribution within the cytoplasm and nucleus of donor (DN) and keratoconus (KC) fibroblasts. Scale bar 100µm.

**Supplementary Figure 4**: Keratoconus study questionnaire.
The patients recruited for the tear sample and in vitro studies were enrolled under the study protocol number 18-00083 at NYU Grossman School of Medicine. GPX3 immunostaining on corneal tissues was performed at the L V Prasad Eye Institute in accordance with the institutional review board (IRB) guidelines under study number LEC-BHR-P-10-23-1114.

#### 18-00083 Keratoconus Genetics Study Questionnaire KCN:

**Research Personnel**

KCN: ______________

Date of Consent:

______________________

Location: _____________

##### Demographic Information

1. Full Name:___________________________
2. Date of Birth: __________________________
3. Email: ________________________
4. Sex
   - ☐ Male
   - ☐ Female
   - ☐ Other
5. Ethnicity
   - ☐ Hispanic or Latino
   - ☐ Not Hispanic or Latino
6. Race
   - ☐ White
   - ☐ Black or African American
   - ☐ American Indian/Alaskan Native
   - ☐ Native Hawaiian or Pacific Islander
   - ☐ Asian
   - ☐ More than one race
   - ☐ Unknown

Keratoconus genetics study
18-00083
Page 1 of 4
Version 1.3
10/30/2020

NYU School of Medicine

#### 18-00083 Keratoconus Genetics Study Questionnaire KCN:

##### Ocular History

7. Do you have a family history of keratoconus?
   - ☐ Yes
   - ☐ No

   If yes, please indicate their relationship to you and their approximate age at diagnosis:

   ______________________________________________________________

8. Have you received a cornea transplant?
   - ☐ Yes
   - ☐ No

   If yes, indicate eye and date:

   __________________________________________________________

9. Have you received any other ocular diagnoses or procedures?
   - ☐ Yes
   - ☐ No

   If yes, describe and date:

   __________________________________________

10. Have you undergone collagen cross-linking?
    - ☐ Yes
    - ☐ No

    If yes, indicate eye and date of procedure:

    _________________________________________________

### 18-00083 Keratoconus Genetics Study Questionnaire  KCN:

11. Do you have a history of any of the following? Check all that apply:
   - ☐ Atopy
   - ☐ Allergy
   - ☐ Eczema
   - ☐ Hay Fever
   - ☐ Asthma
   - ☐ None

12. Are you a regular contact wearer?
   - ☐ Yes
   - ☐ No

   If yes, indicate eye and contact type: ______________________________

13. Have you been diagnosed with keratoconus?
   - ☐ Yes
   - ☐ No

   If yes, please indicated the approximate date of diagnosis:

   Right eye: __________________________________________

   Left eye: ___________________________________________

### 18-00083 Keratoconus Genetics Study Questionnaire KCN:

**Research Personnel Only**

14. MRN: ________________

15. Pentacam
- ☐ Left eye (OS) Km: _____ Astig: ________ Thickness: ______ Kmax: ______
- ☐ Right eye (OD) Km: _____ Astig: ________ Thickness: ______ Kmax: ______

16. Uncorrected visual acuity (sc)
- ☐ Left eye: ______________
- ☐ Right eye:________________

17. Best corrected visual acuity (cc)
- ☐ Left eye: ______________
- ☐ Right eye:________________

18. Date of blood draw: ________________

19. Date of tissue collection: ________________

______________________________________ ___________________
Signature of Research Staff Date
